# Multi-way methods for understanding longitudinal intervention effects on bacterial communities

**DOI:** 10.1101/363630

**Authors:** Ingrid Måge, Christina Steppeler, Ingunn Berget, Jan Erik Paulsen, Ida Rud

## Abstract

**Background:** This paper presents a strategy for statistical analysis and interpretation of longitudinal intervention effects on bacterial communities. Data from such experiments often suffers from small sample size, high degree of irrelevant variation, and missing data points. Our strategy is a combination of multi-way decomposition methods, multivariate ANOVA, multi-block regression, hierarchical clustering and phylogenetic network graphs. The aim is to provide answers to relevant research questions, which are both *statistically valid* and *easy to interpret*.

**Results:** The strategy is illustrated by analysing an intervention design where two mice groups were subjected to a treatment that caused inflammation in the intestines. Total microbiota in fecal samples was analysed at five time points, and the clinical end point was the load of colon cancer lesions. By using different combinations of the aforementioned methods, we were able to show that:

- The treatment had a significant effect on the microbiota, and we have identified clusters of bacteria groups with different time trajectories.
- Individual differences in the initial microbiota had a large effect on the load of tumors, but not on the formation of early-stage lesions (flat ACFs).
- The treatment resulted in an increase in *Bacteroidaceae*, *Prevotellaceae* and *Paraprevotellaceae*, and this increase could be associated with the formation of cancer lesions.

**Conclusion:** The results show that by applying several data analytical methods in combination, we are able to view the system from different angles and thereby answer different research questions. We believe that multiway methods and multivariate ANOVA should be used more frequently in the bioinformatics fields, due to their ability to extract meaningful components from data sets with many collinear variables, few samples and a high degree of noise or irrelevant variation.

## Background

This paper presents a novel strategy for statistical analysis and interpretation of longitudinal intervention effects on bacterial communities. The strategy is based on a combination of multi-way decomposition methods, multivariate ANOVA, multi-block regression, cluster analysis and phylogenetic network graphs. The aim is to produce results that are both *statistically valid* and *easy to interpret.* Although none of the methods are new per se, the novelty lies in the combined application which illustrates their usefulness. Also, two of the methods (PARAFASCA and SO-NPLS) do not have any published applications aside from examples in the original publications [1], [2].

The strategy is illustrated by analysing an intervention design on total microbiota. Such data are multivariate and longitudinal (i.e. three-dimensional) by nature, with samples, bacterial groups and time as the three ways. The data set also has a number of statistical challenges that are often encountered in these types of studies, such as small sample size, many more variables than samples, high degree of irrelevant variation, and missing data points.

A multitude of statistical methods for analysing temporal effects on microbiota have been suggested, but most of them analyse each variable separately in a univariate fashion[3], [4]. The approach presented in this paper is multivariate, meaning that it analyses the total covariance pattern of all bacterial groups (variables) simultaneously. With today’s high-throughput sequencing techniques, the number of bacterial groups can range from hundreds to thousands and univariate analysis is no longer feasible.

Longitudinal measurements of the microbiota give rise to multi-way data. For instance, when the bacterial *composition* is followed over *time* for a group of *subjects*, the data can be arranged in a three-dimensional cube with dimensions *subjects*, *bacterial groups* and *time.* The data cube is often unfolded into a two-dimensional matrix and analysed by standard multivariate methods such as principal component analysis (PCA), principal correlation analysis (PCoA [5]) and partial least squares regression (PLSR [6]). However, this kind of unfolding can be unfavourable for many reasons: First of all, unfolded models are very complex (many parameters need to be estimated), which increases the risk of overfitting and complicates interpretation. Secondly, the information from the unfolded dimensions is intertwined, obscuring the interpretation further. Multiway methods are data decomposition methods that can handle multi-way arrays without unfolding them. Parallel factor analysis (PARAFAC), Tucker and multiway partial squares (N-PLS) have been used since the 1990’s in the fields of psychometrics and chemometrics [7], [8], but we have so far seen few applications in microbiology. Of the few applications that has been reported, three-way decomposition methods have been used to analyse a longitudinal study of the faecal microbiota in pregnant mothers and their children up to two years of age [9], [10]. The usefulness of multiway methods has also been advocated in the field of metabolomics, which share many of the data characteristics of bacterial community data [11], [12].

In intervention studies, the hypothesis is usually that a certain treatment induces a change in the microbiota, which again may lead to changes in phenotype (often some health related parameter). The main research questions are typically:

1. How does the intervention design affect the total microbiota?
2. How do the changes in microbiota affect the subject’s health?
3. What are the individual differences in microbiota between subjects? Are these differences important for the health?

In this paper we propose different variants of multivariate and multiway methods for answering the three main research questions, combined with a number of additional tools for interpreting and visualizing the results. In addition, we use *hierarchical clustering* to identify groups of bacteria that have similar time trajectory responses, and *network graphs* for visualising and interpreting the results in view of the bacteria’s genetic similarities. An advantage of the network graph is that it can be used interactively to combine and explore the results from several models together.

## Methods

An overview of our data analysis strategy is illustrated in Figure 1. First, explorative analysis is performed to get an overview of the data and check for outliers. In this step, the missing data points can also be estimated. Multivariate ANOVA is used to relate the microbiota to the intervention design, addressing research question 1) in the introduction. Multiway regression is used to relate the microbiota to cancer lesions. Specifically, the sequential regression method SO-NPLS is used to separate individual differences from treatment effects in the regression analysis, addressing research question 2) and 3) in the introduction. All these methods are coupled with variable selection procedures that identifies the important bacteria groups. The cluster analysis then identifies bacteria with similar time trajectories, which aid interpretation of the ANOVA and regression models. The resulting model parameters and clusters are finally visualized in a phylogenetic network graph in order to facilitate biological interpretation. All data analysis was done in MATLAB (version R2016a, The Mathworks, Inc.), and the network graph was drawn with Gephi [13].

**Figure 1.**
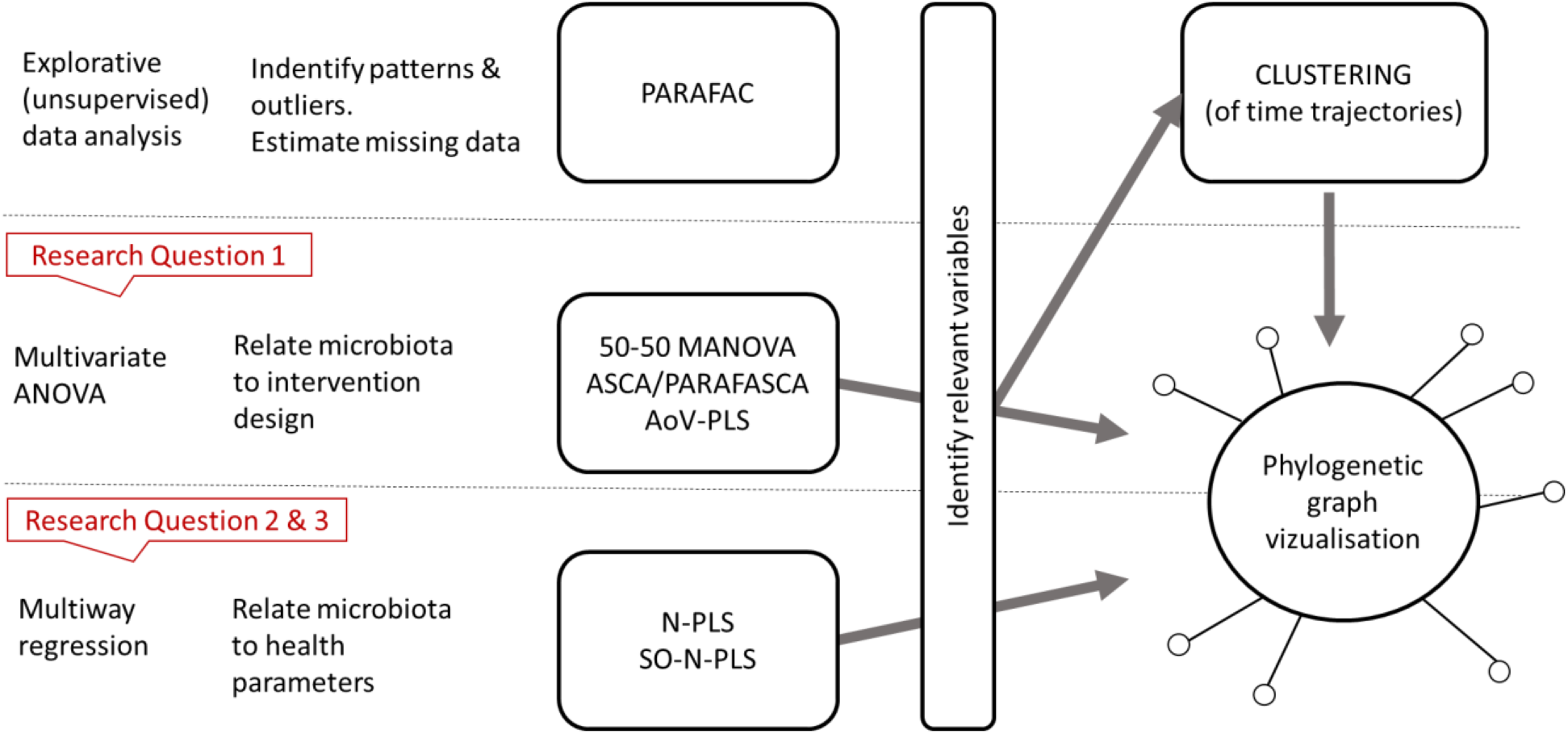
Schematic illustration of the strategy for data analysis and interpretation. First, explorative analysis is performed to get an overview of the data and check for outliers. Then, multivariate ANOVA is used to assess effects of the intervention design, and multiway regression to correlate the microbiota to health parameters. Variable selection, clustering and phylogenetic network graphs are used for biological interpretation of the results.

### Example data set

The data set is from an intervention study on A/J Min/+ mice, which serve as a model system for colorectal cancer. The objective of the study was to investigate how the gut microbiota and formation of cancer lesions is affected by inflammation. A detailed description of the study with biological interpretations is given in [14]. The intervention included sixteen mice, of which eight were treated with dextran sodium sulphate (DSS) via drinking water for 4 days. This treatment induced an inflammation in the intestines. Two groups of mice were used, either twelve (Gr1) or ten (Gr2) weeks old, and these were distributed as evenly as possible between the treatment and control group. Faeces samples were collected from all mice at day 1 of the experiment (before treatment) and at day 5, 8, 15 and 24 (after treatment). The faeces samples were analysed with regard to bacterial composition. At day 24, the mice were sacrificed, and tumorigenesis was quantified by measuring the loads (total areal) of both early-stage lesions (flat ACFs) and tumours. Several other measurements were also done, such as quantification of short-chain fatty acids (SCFA) in faeces and immunobiological analysis of the spleen and Peyer’s patches. These are not used here, but included in reference [14].

The bacterial community was analysed by 16S rRNA amplicon sequencing on a MiSeq (Illumina) sequencer. The reads were processed according to the pipelines in Quantitative Insight Into Microbial Ecology (QIIME) v.1.8 [15]. This pipeline results in a list of Operational Taxonomic Units (OTUs), each representing a phylotype that may be a representative of a bacterial species. The abundance of a specific OTU in a sample is calculated as the number of sequences matching that OTU, relative to the total number of sequences in the sample. Many of the OTUs have zero or very low abundance in a majority of the samples. These were filtered out by applying the criterion that every OTU should have abundance >0.05% in more than half of the samples from at least one of the four treatment/group combinations. The resulting data set consisted of 526 OTUs.

Three faeces samples were missing, from three different animals in the control group at day one, five and eight respectively. The full data set analysed in this paper is illustrated in Figure 2.

**Figure 2.**
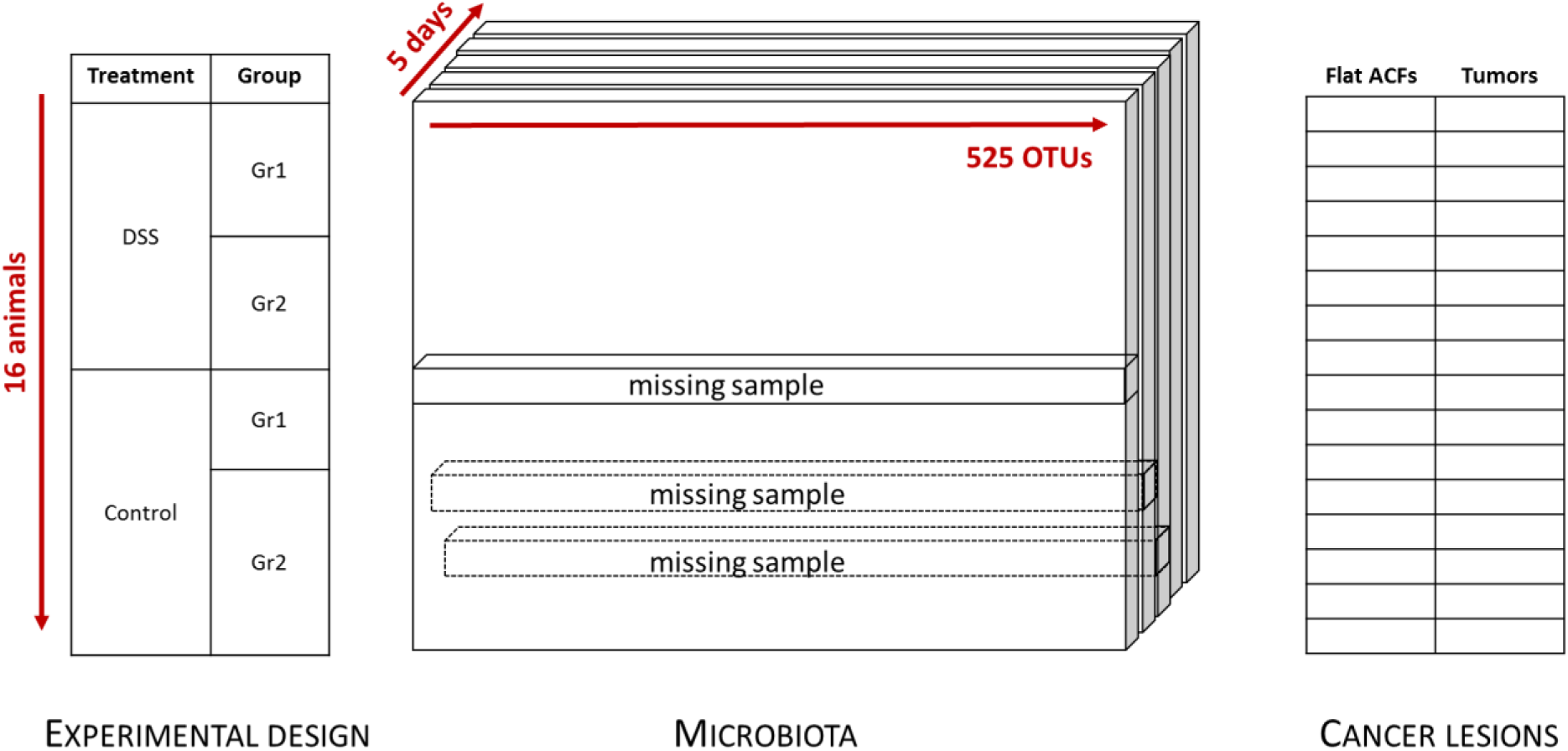
Illustration of data set consisting of the intervention design with sixteen animals, microbiota measured at five occasions (day 1,5,8,15 and 24) and cancer lesions at the end of the study (day 24).

### Multiway data arrays

Multiway data occur when several sets of variables are measured in a crossed fashion, for instance when the microbial *composition* is followed over *time* for a group of *subjects.* This results in a three-way array, also called tensor, with the dimensions (*I* subjects) × (*J* bacterial groups) × (*K* time points). The dimensions are often called *modes.* In the following we refer to scalars as lower-case italics (e.g. *x*), vectors as lower-case bold (e.g. **x**), matrices as uppercase bold (e.g. **X**), and 3-dimensional tensors as underlined uppercase bold (**<u>X</u>**).

Centring and scaling three-way arrays is slightly more complicated than in the two-way case, especially if more than one mode is to be scaled. In general, it is unproblematic to scale within one mode and centre across another mode as long as the centring is done first [16]. For bacterial community data, it is natural to centre across subjects and scale the bacterial groups, to remove relative differences in abundance and variation. The subject mode is centred by unfolding the array into an *I* × *JK* matrix and subtracting the mean of every column. Scaling of variables should be done “slice”-wise, not column-wise. This means that all columns containing a variable should be scaled by the same factor. Each of the *J* variable slices (*I* × *K* matrices) were in this case scaled to unit squared variance by:

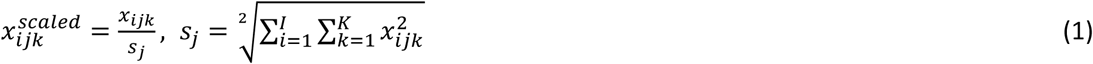

### Explorative analysis using PARAFAC

All data analysis starts with an explorative phase, where the objective is to get a first overview of the data and identify errors, oddities or outlying observations. Principal Component Analysis (PCA) is commonly used for exploration of multivariate data, and PARAFAC is the generalisation of PCA for higher order arrays [17]. It decomposes the data into a few underlying components that summarize the main systematic variation in the data. This means that the main variation can be summarised in simple two-dimensional plots, making the method well suited for explorative analyses of multiway arrays.

A PARAFAC model with *F* components, can be written as a Kronecker product:

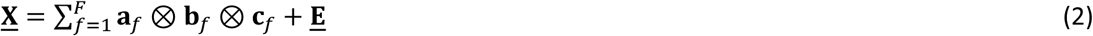

where **a**_*f*_, **b**_*f*_ and **c**_*f*_ are the loadings for component *f* in mode one, two and three respectively, and **<u>E</u>** contains the residuals. The components are defined so that the sum of the squared elements in **<u>E</u>** is minimised. A graphical illustration of a two-component model is given in Figure 3. In contrary to PCA, the PARAFAC components are not necessarily orthogonal. This is important to keep in mind when interpreting the loadings. Several constraints may be imposed on the loadings for one or several modes, such as non-negativity, unimodality and orthogonality. This is often based on knowledge about how the true underlying components should behave. In this work constraints are not applied.

**Figure 3.**
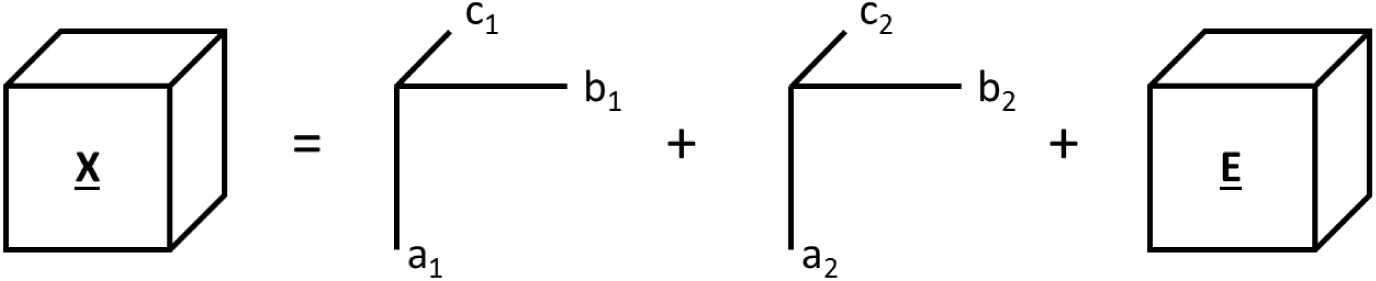
*Graphical illustration of a two*-*component PARAFAC model. The three*-*dimensional data array* **<u>X</u>** *is decomposed into two sets of loadings (a*, *b and c) corresponding to the three dimensions*, *and a residual array* **<u>E</u>**. *The mathematical formulation is given in Equation (2)*.

The PARAFAC model is fitted using an alternating least squares algorithm, which has the advantage of handling missing values. Missing values are estimated by multiplying the loadings (as in Equation 2) in each iteration, and the algorithm continues until convergence of the missing values and the overall model fit. In this work we have used the single imputation (expectation maximisation) algorithm, which is shown to give good results for up to 60% randomly missing data [18].

The number of components can be selected by inspecting the *core consistency diagnostic* as a function of number of components, which should be high (80-100%) for a valid model. See [17] for details. An implementation of PARAFAC can be found in the N-way toolbox for MATLAB [19].

### Relating microbiota to the intervention design

Analysis of Variance (ANOVA) is the appropriate method for analysing data from designed experiments with one response variable at the time, but it is not suited for analysing many collinear responses. Alternative methods for multiple collinear responses are for instance ANOVA-Simultaneous Component Analysis (ASCA) [20], AoV-PLS [21] and fifty-fifty MANOVA [22]. ASCA is a method that combines ANOVA with the dimension reduction and interpretation tools of PCA. The basis of ASCA is the same linear model as used in ANOVA, splitting the total variation into contributions of design factors and their interactions. Here we analyse an experimental design with the three factors mouse group (*Gr*), treatment (*Trt*) and *Time.* The data cube is unfolded by stacking the five time points on top of each other, giving the matrix **X**_*unf*_ with dimensions *IJ* × *K*. The data matrix is then split into submatrices according to the model:

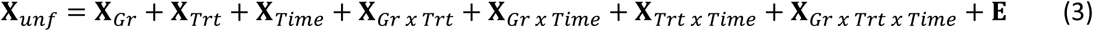

Each of the effects can be interpreted by decomposing the sub-matrices using PCA, and the statistical significance can be evaluated by permutation testing. When the data comes from an unfolded multidimensional array, the effects that contain more than one dimensions can be refolded and decomposed by PARAFAC instead of PCA, yielding PARAFASCA [1]. Here, the effects involving *Time* are refolded into a three-dimensional cube and interpreted by PARAFAC.

AoV-PLS is very similar to ASCA. The main difference lies in the interpretation and significance testing of each submatrix in Equation (3), which is done by Partial Least Squares (PLS) regression [6] instead of PCA. PLS regression is commonly used to relate multiple collinear variables to one or several responses. In AoV-PLS, a PLS regression model is fitted using **X**_*Effect*_ as response and (**X**_*Effect*_ + **E**) as regressor. If the regression model is statistically significant, the effect is considered as significant. AoV-PLS is a newer and less established method than ASCA, and a thorough comparison of the two methods has not been done yet.

In fifty-fifty MANOVA [22], [23], the dimensionality of the data is reduced by PCA and a so-called “fifty-fifty F-test” is used to assess the significance of each effect. The fifty-fifty MANOVA results in a table resembling the traditional ANOVA table for univariate responses and can be interpreted in the same way, but do not offer visualization tools for displaying the effects.

### Relating microbiota to tumorigenesis

As mentioned in the previous section, PLS regression can be used to relate multiple collinear variables to one or several responses [6]. The original PLS regression handles two-way matrices only, but an extension that handles multiway explanatory variables exists (N-PLS, [24]). Other extensions of PLS regression handle multiple blocks of explanatory variables, for instance multiblock PLS [25] and Sequential Orthogonalised PLS (SO-PLS, [26]). Here, SO-PLS and N-PLS is combined as described in [27] by using N-PLS instead of PLS in the sequential modelling. This is done in order to analyse the effect of the initial bacterial composition (before intervention) and the *incremental* effect of the intervention. The data arrays used in this particular SO-NPLS model are illustrated in Figure 4. The first block is the initial bacterial composition, denoted ***X**_Init_*, and the second block is the bacterial composition for the different time points after the treatment and is denoted (***X**_Trt_*). The predictive part of the *X_Init_* can be interpreted as the influence of the group and individual differences on cancer development. The predictive part of ***X**_Trt_* can be interpreted as the additional effect caused by the intervention-induced changes in microbiota. The number of predictive components for each block (*a_1_* and *a_2_*) is usually determined by cross-validating all combinations of *a_1_* and *a_2_* up to a maximum number, selecting the combination that balances model fit and parsimony. Details on the algorithm can be found in [2].

**Figure 4.**
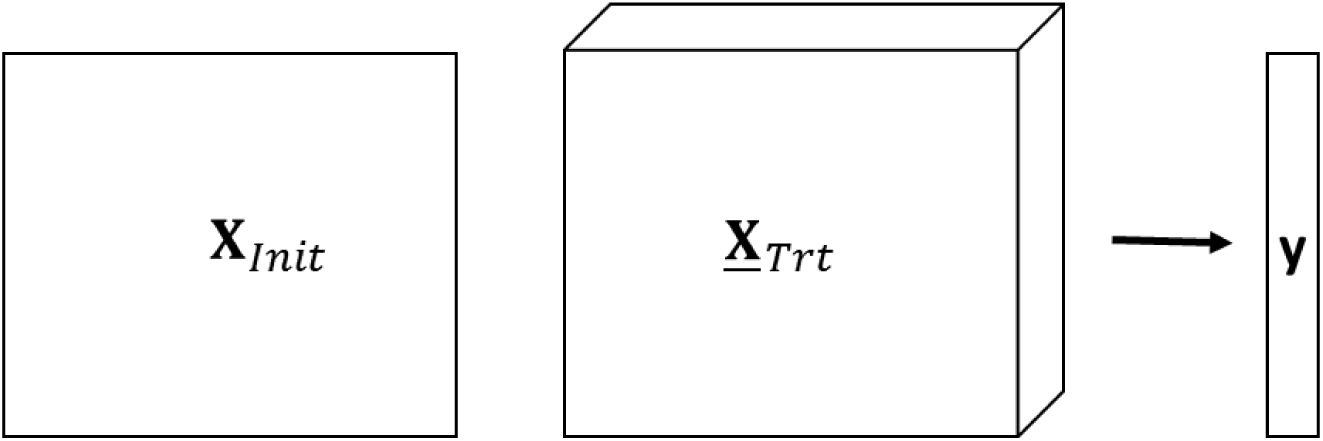
*Data matrices in the SO*-*NPLS model. The first data matrix (**X**_Inlt_) contains the initial microbiota at day 1*, *i.e. before intervention. The second data cube* (***<u>X</u>**_Trt_*) *contains the microbiota at day 1*, *5*, *8*, *15 and 24. The response (***y***) is the load of either early*-*stage lesions or tumours*.

### Identifying important variables

For biological interpretation, it is crucial to identify the bacteria groups that are associated with an effect or component in any of the above-mentioned models. In the framework of fifty-fifty MANOVA, a rotation test can be used to calculate p-values that are adjusted for multiple comparisons [27], [28]. The adjusted p-values control the false discovery rate. In ASCA, the magnitude of the loadings in the OTU mode gives an indication of which OTUs that are related to each model component. This method is however not pursued here, as it is hard to select the cut-off value. An advantage of AoV-PLS over ASCA is that any variable selection method suitable for PLS regression may be applied. There are a number of such variable selection methods [29], and a comparison of these is not in the scope of this paper. We have chosen to use the well-established method Variable Importance in Prediction (VIP, [30], [31]), and for SO-NPLS we use the extension of VIP for multiway regression [32].

### Clustering of time trajectories

To facilitate interpretation, the OTU’s time trajectories were grouped by hierarchical clustering using Euclidean distances and Ward linkage. The clustering was only performed on the OTUs that were identified as significantly affected by the DSS treatment. Clustering was performed on the time trajectories averaged over mice in the treatment group.

The number of clusters was determined by visual inspection of the dendrograms, in combination with the cluster validation techniques silhouette plots [33], within-between statistics (WB) and the Hubert-Gamma statistics (HG) [34]. In addition, the stability of clusters were assessed by first classifying OTUs for each individual mouse to the clusters identified from the average over all mice, using quadratic discriminant analysis (QDA) [35]. Then, the number of times the OTUs were classified to each cluster was counted. The stability of the partition was assessed by looking at the confusion matrix between classification into the cluster with maximum number of counts, and the cluster obtained for the average.

### Phylogenetic network graphs

Phylogenetic relations between the bacteria groups can be visualized in a force-directed network graph, where the bacteria groups are represented by nodes and their genetic similarities by edges between nodes. The purpose is to position the nodes in a two-dimensional space so that similar nodes are grouped close together, while dissimilar nodes are positioned far apart. A number of different algorithms exist for calculating the graph layout. Most of them seek to find a minimum-energy equilibrium state of the system given various physical laws and restrictions. We have chosen to use the ForceAtlas2 method within the Gephi software [13], [36]. This is a fast method that also takes the edge weights (degree of similarity) into account when drawing the graph. In this work, each OTU was represented by a 16s DNA sequence of length ≈250, and the genetic distances between all pairs of bacteria groups were calculated as the proportion of sites at which the two sequences are different (p-distance, [37]). The p-distance is close to one for poorly related sequences and approaching zero for similar sequences. The edge weights were then defined as one minus the p-distance.

## Results

In this section we exemplify how the methods can be used to answer the three research questions given in the introduction. We start by exploring the composition of the microbiota through a phylogenetic network graph. Then we check the data and estimate missing values using PARAFAC. We assess the effects of the experimental factors (research question 1) by multivariate ANOVA, including the three-way interpretation method PARAFASCA. Finally, we relate the microbiota to formation of cancer lesions, splitting individual differences and treatment effects using SO-NPLS regression (research question 2 and 3).

### Exploring the microbial composition

Figure 4a) shows the phylogenetic network graph, coloured by taxonomic family. This graph illustrates the genetic relationship between the 526 OTUs. The most dominating families are *S24*-*7*, *Bacteroidaceae* and *Paraprevotellaceae* from the *Bacteroidetes* phylum, in addition to *Ruminococcaceae* and *Lachnospiraceae* from the *Firmicutes* phylum.

### Exploring the data structure and estimating missing values

The three-way microbiota array (Figure 2) was decomposed by unconstrained PARAFAC, and a three-component model was selected by examining the residual variance and core consistency (which was 93%). Two of the components showed a clear treatment effect, while the third component distinguished between the two mouse groups. The three components explain 27% of the total variation in microbiota, meaning that 73% of the variation is due to individual differences between subjects and random variation. This is expected for this type of data, and the model was deemed suitable for estimating the three missing samples by multiplying the loadings according to Equation 2. The missing data were substituted by these estimates in all subsequent analysis. More details about the PARAFAC model is not shown, as it is more meaningful to interpret the data by means of multivariate ANOVA and regression.

### Relating microbiota to the intervention design

The PARAFAC indicated clear effects of group and treatment, but a more formalized decomposition of the effects is obtained by ASCA (Equation 3) or the equivalent AoV-PLS and fifty-fifty MANOVA models. Explained variances and p-values from ASCA and fifty-fifty MANOVA are reported in Table 1. AoV-PLS gives the same effect sizes as ASCA, and is therefore not included in the table. ASCA and fifty-fifty MANOVA give almost identical results with regard to effect sizes and statistical significance. The largest effect on microbiota is caused by the DSS treatment, Time and their interaction, as expected. There is also a statistically significant (although small) difference between groups, and an even smaller interaction between group and treatment.

**Table 1.** Analysis of variance results for the microbiota (using both ASCA and fifty-fifty MANOVA) and tumorigenesis

| Factor | Microbiota |  |  |  | Load of flat ACFs |  | Load of tumors |  |
| --- | --- | --- | --- | --- | --- | --- | --- | --- |
|  | ASCA |  | fifty-fifty MANOVA |  | univariate ANOVA |  | univariate ANOVA |  |
|  | Effect size (explained variance) (%) | p-value* | Effect size (explained variance) (%) | p-value+ | Effect size (explained variance) (%) | p-value | Effect size (explained variance) (%) | p-value |
| Gr | **4.2** | **<0.001** | **4.2** | **<0.001** | 4.9 | 0.159 | 8.6 | 0.208 |
| Trt | **10.4** | **<0.001** | **10.2** | **<0.001** | **55.0** | **<0.001** | 6.4 | 0.276 |
| Time | **11.4** | **<0.001** | **11.5** | **<0.001** |  |  |  |  |
| Gr x Trt | **2.2** | **<0.001** | **2.3** | **<0.001** | **9.9** | **0.054** | **24.1** | **0.046** |
| Gr x Time | 3.5 | 0.756 | 3.5 | 0.104 |  |  |  |  |
| Trt x Time | **7.9** | **<0.001** | **7.7** | **<0.001** |  |  |  |  |
| Grp x Trt x Time | 3.1 | 0.966 | 3.1 | 0.879 |  |  |  |  |
| Residuals | 57.4 |  | 57.3 |  | 26.0 |  | 58.4 |  |

\* Based on permutation testing + Based on the fifty-fifty F-test.

Generally, main effects should not be interpreted alone when there is a significant interaction effect. The effect matrices for Trt, Time and Trt × Time where therefore The important OTUs for each effect was evaluated by both VIP (using AoV-PLS) and rotation tests, as described in the methods section. The VIP threshold was set to one. Venn diagrams in Figure 6 illustrate the agreement between the two methods, together with the correlation between variable ranking from VIP- and p-values (spearman’s rho). The agreement is very good for the combined effect of *Treatment* and *Treatment × Time*, which is also by far the largest effect. The agreement is lower for the smaller effects of *Group* and *Group × Treatment*, where the rotation test is much more conservative than VIP.

**Figure 5.**
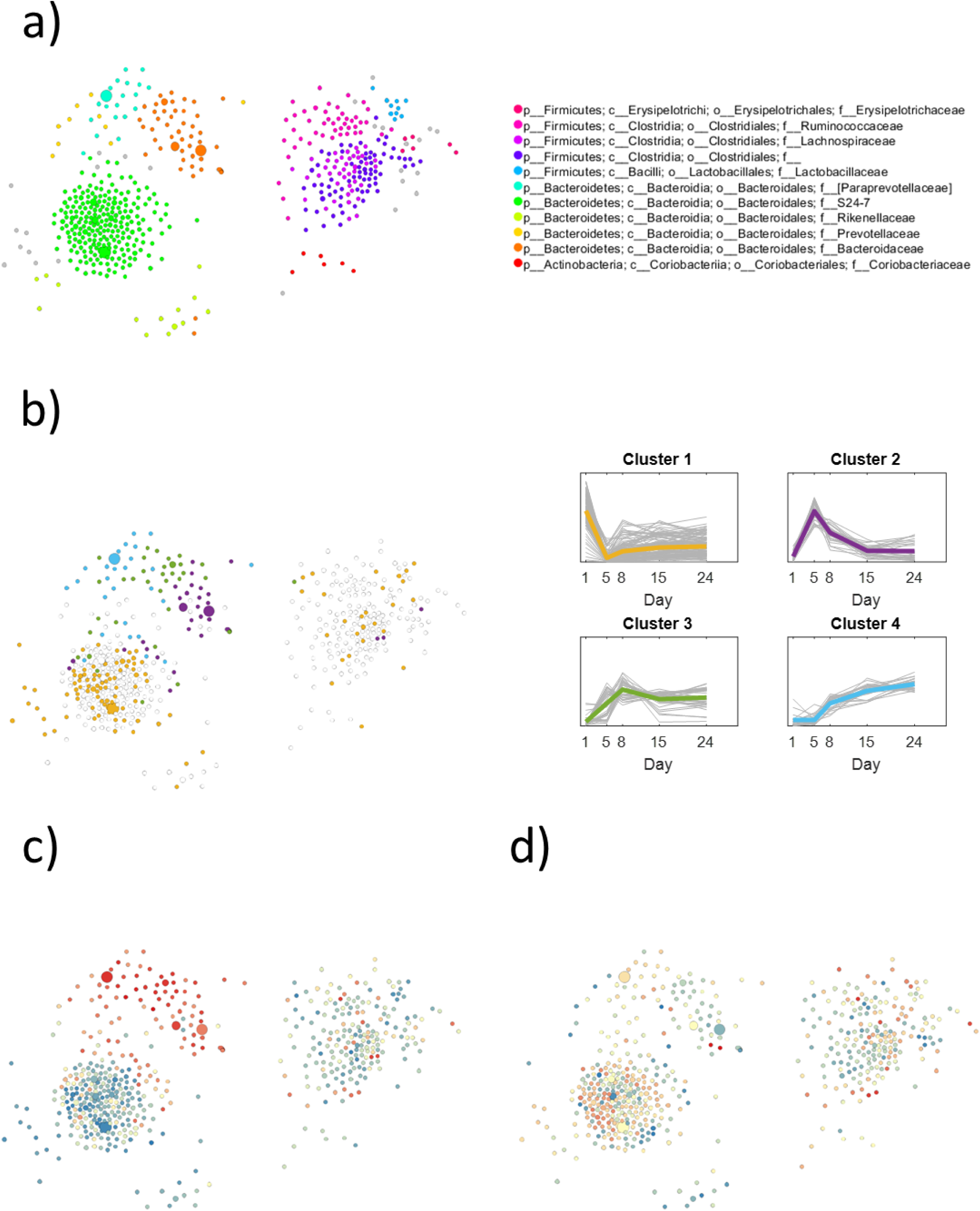
Network graph illustrating the genetic similarities between the 526 OTUs. a) coloured by phylogenetic family, b) coloured by time trajectory cluster, c) coloured by loadings from flat ACF regression model (component 1), d) coloured by loadings from flat ACF regression model (component 2), colour gradient from blue (negative) through white (zero/unimportant) to red (positive). Subplot c) and d) correspond to the scores and loadings in Figure 8..

**Figure 6.**
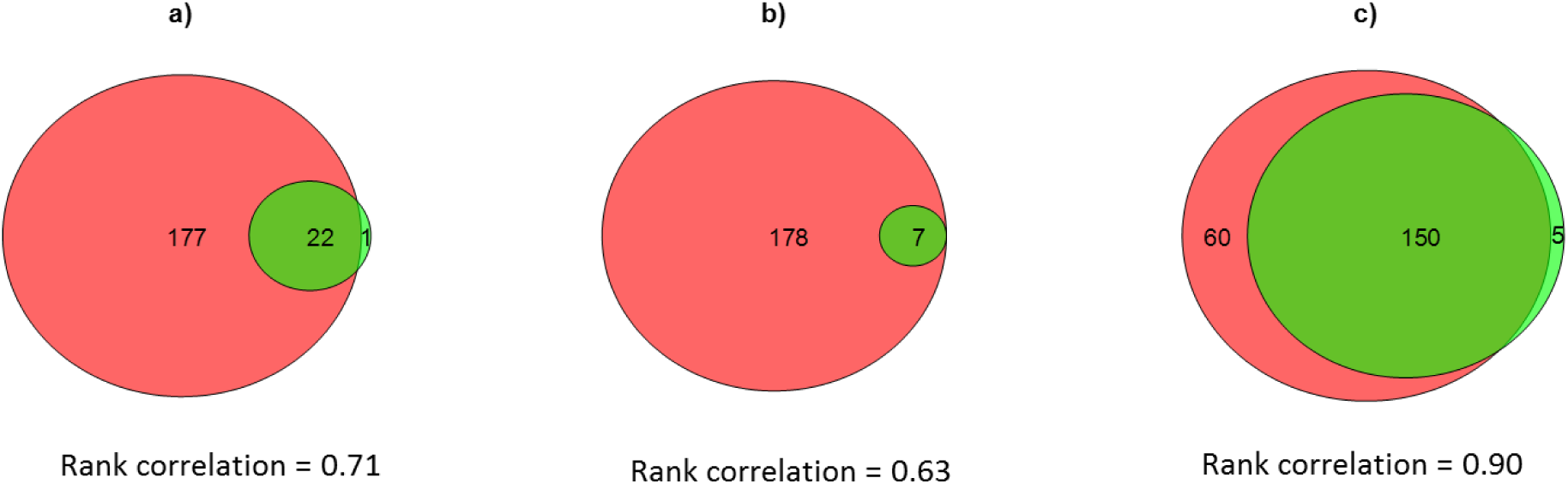
Venn diagrams showing correspondence between important variables from AoV-PLS combined with VIP (pink/left) and rotation tests (green/right) for a) Group effect, b) Group × Treatment interaction and c) (Treatment + Treatment × Time) effects. A cutoff value of one was used for VIP, and 5% significance level for the rotation tests.

In ASCA and AoV-PLS, the effects can be interpreted by PCA (or PLS regression) on the effect matrices, as described in the methods section. The *Group* and *Group × Treatment* effects have one degree of freedom each and are therefore described by one principal component each. PCA scores and loadings for these effects are shown in Figure 7a-b. The OTU loadings are represented by dots which sizes correspond to abundance, and those found to be significant by either rotation tests or VIP are filled with black colour. Note that most of the significant OTUs for these effects are small, i.e. in low abundance.

**Figure 7.**
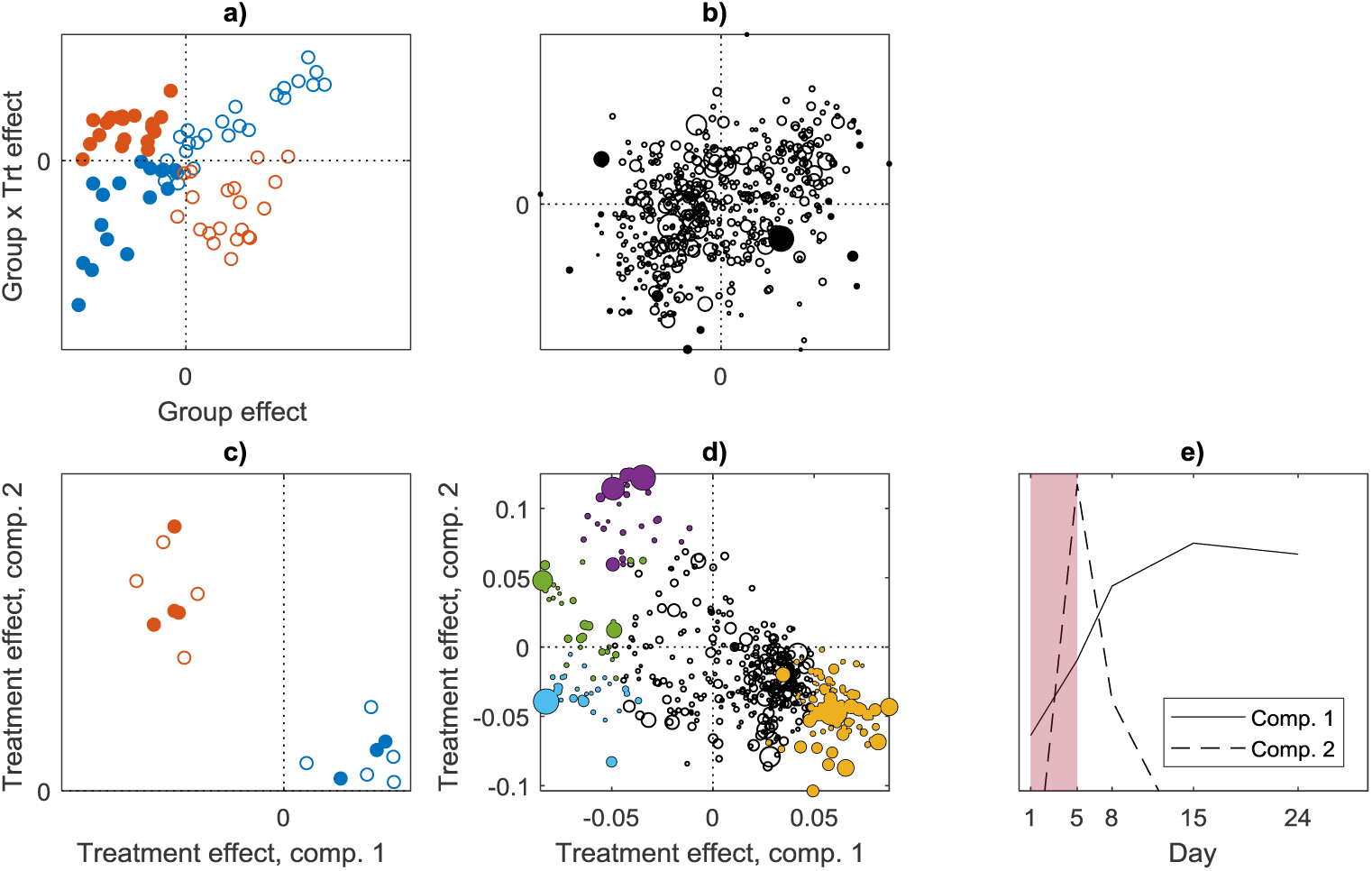
*ASCA scores (a) and OTU loadings (b) for* group *and* group × treatment *effects*, *and PARAFASCA scores (c)*, *OTU loadings (d) and time loadings (e) on* (Treatment + Treatment × Time) *effects. Sample scores in a) and c) are coded according to treatment (red* = *DSS*, *blue* = *control) and mouse group (filled circles* = *Group 1*, *open circles* = *group 2). Filled OTU loadings are those found to be significant by either rotation tests or VIP*, *and the colours in d) correspond to time trajectory clusters, equivalent to those in Figure 5. The shaded region in e) represent the treatment period*.

There is a significant interaction effect between *Time* and *Treatment*, which means that the main effects should not be interpreted separately. The combined effect matrices of *Treatment*, *Time* and *Treatment × Time* was therefore investigated by PARAFAC (i.e. PARAFASCA), and a two-component model was selected. The PARAFAC scores and loadings are given in Figure 7c-e. The two components explain 42% and 23% of the variation in the combined *Treatment*, *Time* and *Treatment × Time* effect. The time loadings in Figure 7e show that there is one phenomenon (component one) that increases steadily and stays high over time, while the second component represents a rapid increase that quickly converges back towards the starting point. This does not mean that all bacteria follow one of these trajectories, but rather that all the trajectories can be expressed as linear combinations of the two. The OTU loadings in Figure 6d are the coefficients of these linear combinations. The OTU loadings that were found to be significant are coloured according to clusters with similar time trajectories (see next section for details). Note that many of the largest dots are coloured, meaning that many of the highly abundant OTUs are significantly affected by treatment and time.

### Cluster analysis of time trajectories

The 215 OTUs that were found to be significantly affected by treatment either from AoV-PLS/VIP or rotation tests, were further investigated by hierarchical cluster analysis. Inspection of the dendrogram and common cluster validation techniques (silhouettes, WB and HG) indicated two to five clusters. With two clusters, one group of OTUs decreases after treatment, and the other increases. This partition is stable (across mice) as 86% and 98% of OTUs were classified to the same cluster as the average mouse.

When the data are split further, into 3-5 clusters, the increasing OTUs are split into subgroups with different time trajectories. These clusters are less stable, indicating variation between mice in when the increase occurs. Based on cluster validation measures and stability estimations, we selected to split the OTUs into four groups, leading to one decreasing cluster (Cluster 1), and three increasing clusters (Cluster 2-4).

The decreasing cluster (Cluster 1) is very stable and show a steep decrease after treatment. The three increasing clusters represent OTUs that either increase at day five, followed by a decrease down to the base level (Cluster 2), increase at day five or eight, and staying at an elevated level (Cluster 3) or steadily increasing from day eight (Cluster 4). Cluster 2 and 4 are quite stable, while cluster 3 is less stable.

The clusters are visualized in the phylogenetic graph in Figure 5b, revealing that the *S24*-*7* family generally decreases with treatment (Cluster 1), while *Bacteroidaceae* increases quite rapidly (cluster 2 or 3), while *Prevotellaceae* and *Paraprevotellaceae* increase at a later stage (Cluster 4). Families under the *Firmicutes* phylum generally decrease with treatment (Cluster 1).

### Relationship between microbiota and cancer

ANOVA of the two response variables *load of tumours* and *load of flat ACFs* (see Table 1) show that the early-stage tumours (flat ACFs) are highly affected by the treatment. The treatment effect is much smaller for the tumour load, but the differences between mouse groups is more evident. In order to assess the direct relationship between microbiota and cancer, the initial microbiota (**X**_*init*_, a two-dimensional data matrix) and the intervention effect (**<u>X</u>**_*trt*_, a three-dimensional data cube) was correlated to the load of flat ACFs and tumours using the sequential SO-NPLS method. The number of components were selected by comparing the cross-validated fit for all combinations of numbers of components from both blocks. All the reported explained variances are based on full cross-validation.

For early-stage lesions (flat ACF), the optimal model has zero components from **X**_*init*_ and two components from **<u>X</u>**_*trt*_ (see Figure 8a). This means that the individual differences in microbiota before the intervention did not influence the formation of new lesions. The resulting model is therefore a two-component N-PLS model, explaining 64% of the variation in flat ACFs. The scores and loadings are shown in 8b)-e). We see from the score plot (b) that the first component, explaining 56% of the variation in early-stage lesions, can be interpreted as a general treatment effect. This corresponds to the main effect of treatment from the ANOVA (Table 1). The second component, explaining an additional 8%, represents a treatment effect that is specific for *group 1* mice, representing the interaction effect from ANOVA. The time domain loadings (Figure 8d) reveal that the main treatment effect corresponds to a sudden shift in microbiota (at day 5), while the second component is related to whether the increase is stable or converge back towards the starting point. The loadings in the OTU domain are also illustrated in Figure 5c)-d), in order to understand how they relate to the phylogeny. From 5c) we see that the *S24*-*7* family as well as most of the *Firmicutes* phyla are negatively correlated to formation of flat ACFs, while *Bacteroidaceae*, *Prevotellaceae* and *Paraprevotellaceae* are positively correlated to the formation of flat ACFs. From 5d) we see that a subgroup of S24-7 are coloured red, meaning that the negative correlations (seen in component 1) for these OTUs are stronger for the mice in *group 1.* The opposite is seen for *Bacteroidaceae*.

**Figure 8.**
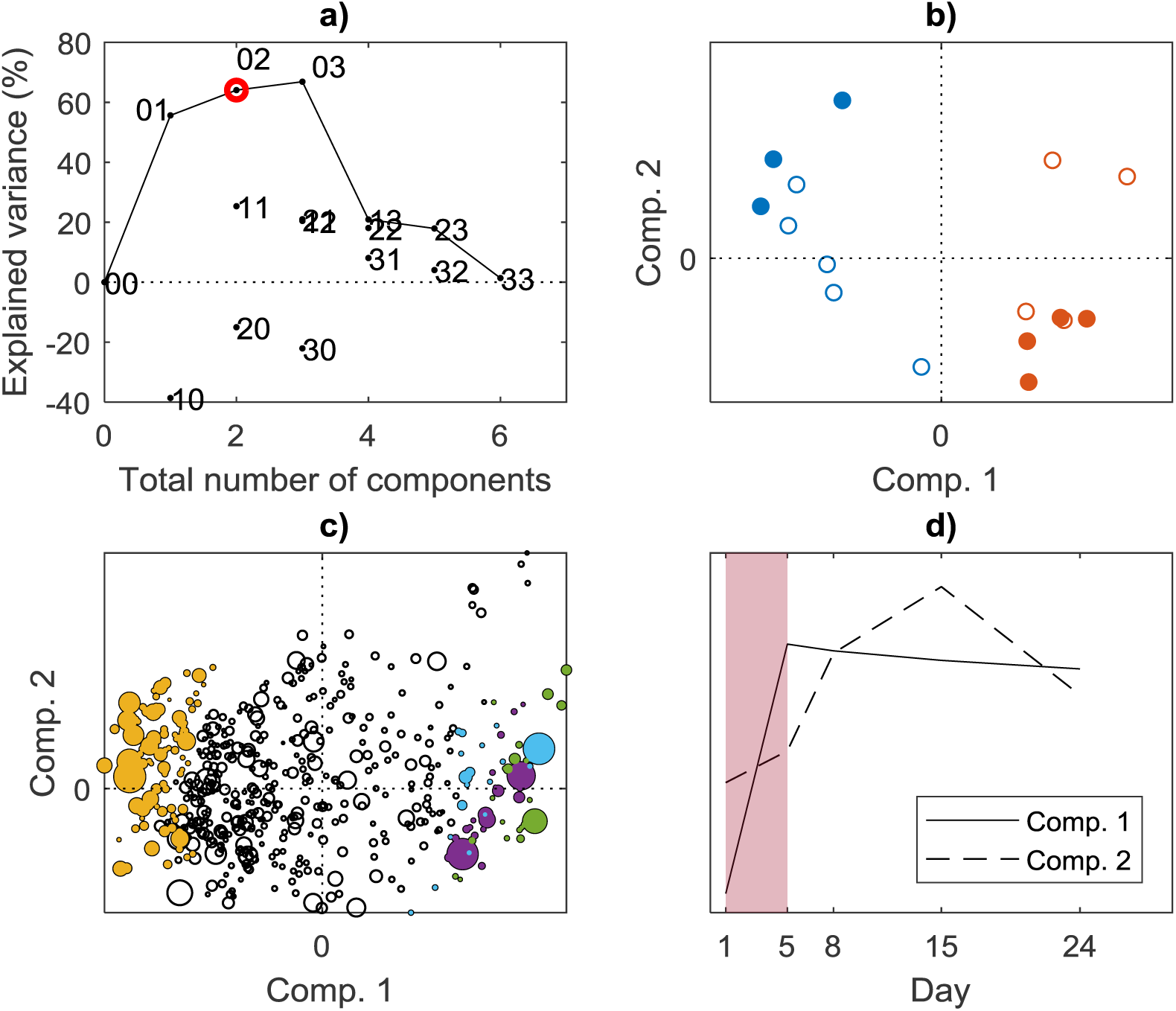
Regression model relating microbiota to early-stage lesions (flat ACFs). a) Mage plot for selecting the number of components in the SO-NPLS model. Each dot represents a candidate model, and the two numeric characters indicate the numbers of components from **X**_init_ and **<u>X</u>**_trt_ respectively. The solid line is the trace of the maximum explained variance. The selected model (marked with a red circle) contains zero components from **X**_init_ and two components from **<u>X</u>**_trt_. b) Subject scores, coded according to treatment (red = DSS, blue = control) and mouse group (filled circles = Group 1, open circles = group 2). c) OTU loadings, the size of each dot represents the abundance. Filled dots are those OTUs found to be significant by VIP, and the colours correspond to time trajectory clusters, equivalent to those in Figure 5. d) Time loadings. The shaded region represents the treatment period.

For explaining the load of tumours, the optimal model has two components from **X**_*init*_ (explaining 34% of the variation in tumour load) and one component from **<u>X</u>**_*trt*_ (explaining and additional 10%), see Figure 9a). It is therefore the individual differences *before treatment* that influence tumour load the most, indicating that these tumours might have been initiated before treatment. The scores and OTU loadings from **X**_*init*_ are given in Figure 9b)-c). The components are not directly related to mouse groups, meaning that there are other individual differences that are related to tumour load. The scores from **<u>X</u>**_*trt*_ (Figure 9d) show that the treatment effect is larger for *group 1* mice, equivalent to what we saw for flat ACFs. The OTU loadings from **<u>X</u>**_*trt*_ are very similar to those from the flat ACFs model, as expected.

**Figure 9.**
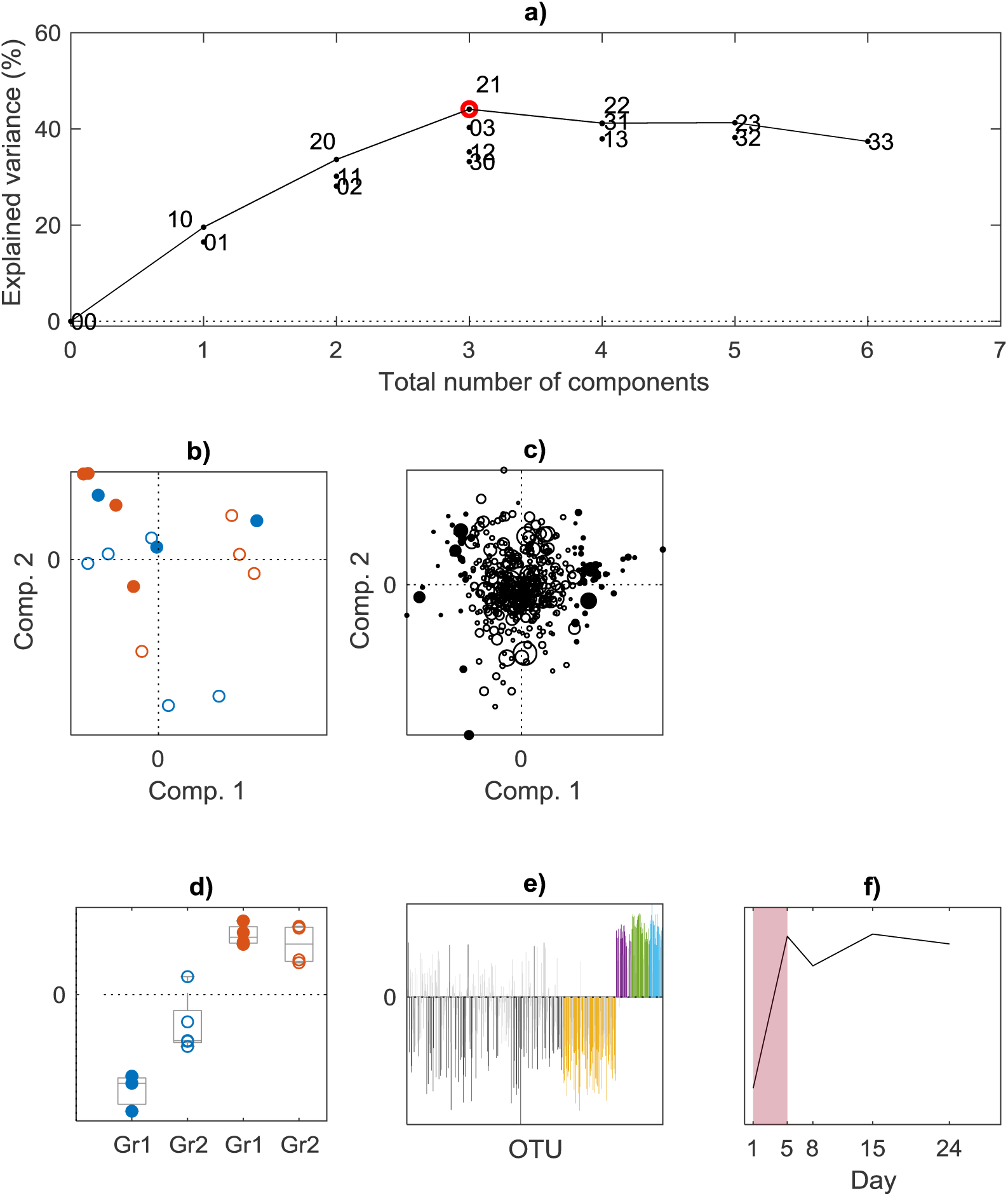
Regression model relating microbiota to load of tumours. a) plot for selecting the number of components in the SO-NPLS model. Each dot represents a candidate model, and the two numeric characters indicate the numbers of components from **X**_init_ and **<u>X</u>**_trt_ respectively. The solid line is the trace of the maximum explained variance. The selected model (marked with a red circle) contains two components from **X**_init_ and one components from **<u>X</u>**_trt_. b) Subject scores from **X**_init_, coded according to treatment (red = DSS, blue = control) and mouse group (filled circles = Group 1, open circles = group 2). c) OTU loadings from **X**_init_. The size of each dot represents the abundance, and filled dots are those OTUs found to be significant by VIP. d) Boxplot of sample scores from the single **<u>X</u>**_trt_ component, with individual scores marked as in b). e) bar plot of OTU loadings from the single **<u>X</u>**_trt_ component, coloured by time trajectory clusters equivalent to Figure 5. d). f) Time loadings from **<u>X</u>**_trt_. The shaded region represents the treatment period.

## Discussion

Even if three-way decomposition methods have existed since the 1970’s, their use in bioinformatics is so far limited. One reason for this might be that they are not well known in the community, even if implementations of PARAFAC and N-PLS are available for both R and MATLAB. Methods such as PARAFASCA and SO-NPLS are on the other hand quite new, and to our knowledge, these methods have not been applied in any scientific field yet. This paper shows that three-way decomposition methods generally have a great potential for analysing and interpreting longitudinal microbial data. One of the main advantages is that such methods allow for graphical displays to summarise large proportions of data, enabling the researchers to get a quick overview of the results. Another advantage is that the solutions are heavily constrained, which implies that they are less prone to overfitting. This is especially important for data with few samples and many variables. A doubt one might have about using such methods for longitudinal data is that they assume trilinearity, meaning that there is a true underlying relationship between the three modes according to Equation (2). This is not necessarily true for these types of data, and more research could be done on the implications of this assumption. However, the example in this paper shows that the three-way methods can extract meaningful components even if these only explain a small part (approx. 25%) of the total variation in the data. From other scientific fields, there are also many examples of successful use of PARAFAC and N-PLS in situations where true trilinearity is not necessarily present, for instance in references [38]–[41].

Multivariate ANOVA by either ASCA, AoV-PLS or fifty-fifty MANOVA consider the underlying covariance between variables instead of one variable at the time. This also prevents overfitting (or false discoveries) when the variables are not independent, as is the case with bacterial strains. Our results show that all methods give very similar results regarding overall effect sizes, but this might not always be the case. A systematic comparison of these three methods has not been done, and this is a topic for further research. The methods for selecting important OTUs give somewhat different results, especially for the factors with small overall effect sizes (*Group* and *Group × Treatment* in our example). This is logical, since there is a higher uncertainty associated with these effects. In addition, a multitude of studies has shown that different variable selection methods give different results, and there is no one method that is always better than others. We therefore recommend to always use several methods and compare the results, to validate the findings. Note that only a few of the OTUs detected as significant with rotation testing were not contained in the sets detected by AoV-PLS/VIP (figure 6).

The experimental design is slightly unbalanced in our case, due to different numbers of mice in each group. We do not expect this to affect the results much in our example, but for more severe unbalance the newly developed ASCA+ method should be applied [42].

## Conclusion

In this paper we have illustrated how multiway methods can be used to analyse and interpret longitudinal microbiome data. We also show how these methods can be combined with phylogenetic network graphs and clustering techniques, for better interpretation of effects.

We have shown that by applying several methods in combination, we are able to “view” the system from different angles and thereby answer different research questions. We believe that multiway methods and multivariate ANOVA should be used more in the bioinformatics fields, due to their ability to extract few, meaningful components from data sets with many collinear variables, few samples and a high degree of noise or irrelevant variation.

## Declarations

### Ethics approval

The experiment was approved by the Norwegian Animal Research Authority and executed in compliance with the local and national regulations associated with animal experiments (application ID: 6906) at the at the experimental animal facility at the Norwegian University of Life Science, Campus Adamstuen.

### Consent for publication

Not applicable.

### Availability of data and material

The dataset supporting the conclusions of this article is available from the authors upon request.

### Author’s contributions

I. Måge analysed the data and wrote the manuscript. C. Steppeler conceived, designed and conducted the mouse experiment and edited the manuscript. I. Berget performed the hierarchical cluster analysis and edited the manuscript. Jan Erik Paulsen conceived and designed the mouse experiment and edited the manuscript. I. Rud was responsible for the microbiota analysis, interpreted the results and edited the manuscript. All authors read and approved the final manuscript.

## Acknowledgements

We like to thank Linn Emilie Knutsen for the assistance during the finalization of the animal work and Merete Rusås Jensen for excellent technical assistance on the analysis of microbiota.

## References

[1] J. J. Jansen, R. Bro, H. C. J. Hoefsloot, F. W. J. Van Den Berg, J. A. Westerhuis, and A. K. Smilde, “PARAFASCA: ASCA combined with PARAFAC for the analysis of metabolic fingerprinting data,” J. Chemom., vol. 22, no. 2, pp. 114–121, 2008.

[2] A. BiancoliMo, T. Næs, R. Bro, and I. Måge, “Extension of SO-PLS to multi-way arrays: SO-N-PLS,” Chemom. Intell. Lab. Syst., vol. 164, pp. 113–126, 2017.

[3] K. Faust, L. Lahti, D. Gonze, W. M. de Vos, and J. Raes, “Metagenomics meets time series analysis: unraveling microbial community dynamics,” Curr. Opin. Microbiol., vol. 25, pp. 56–66, 2015.

[4] X. Zhang, H. Mallick, Z. Tang, L. Zhang, X. Cui, A. K. Benson, and N. Yi, “Negative binomial mixed models for analyzing microbiome count data.”

[5] J. C. Gower, Gower, and J. C., “Principal Coordinates Analysis,” in Wiley StatsRef: Statistics Reference Online, Chichester, UK: John Wiley & Sons, Ltd, 2015, pp. 1–7.

[6] H. Martens and T. Næs, Multivariate Calibration. Chichester: John Wiley & Sons, Ltd, 1989.

[7] P. M. Kroonenberg, “My Multiway Analysis: From Jan de Leeuw to TWPack and Back,” J. Stat. Softw., vol. 73, no. 3, pp. 1–22, 2016.

[8] A. K. Smilde, R. Bro, and P. Geladi, Multi-way analysis: Applications in the chemical sciences. J. Wiley, 2004.

[9] A. Ravi, E. Avershina, S. L. Foley, J. Ludvigsen, O. Storrø, T. Øien, R. Johnsen, A. L. McCartney, T. M. L’Abée-Lund, and K. Rudi, “The commensal infant gut meta-mobilome as a potential reservoir for persistent multidrug resistance integrons,” Sci. Rep., vol. 5, p. 15317, Oct. 2015.

[10] E. Avershina, O. Storrø, T. Øien, R. Johnsen, P. Pope, and K. Rudi, “Major faecal microbiota shifts in composition and diversity with age in a geographically restricted cohort of mothers and their children,” FEMS Microbiol. Ecol., vol. 87, no. 1, pp. 280–290, Jan. 2014.

[11] A. K. Smilde, J. A. Westerhuis, H. C. J. Hoefsloot, S. Bijlsma, C. M. Rubingh, D. J. Vis, R. H. Jellema, H. Pijl, F. Roelfsema, and J. van der Greef, “Dynamic metabolomic data analysis: a tutorial review.,” Metabolomics, vol. 6, no. 1, pp. 3–17, Mar. 2010.

[12] M. M. W. B. Hendriks, F. A. van Eeuwijk, R. H. Jellema, J. A. Westerhuis, T. H. Reijmers, H. C. J. Hoefsloot, and A. K. Smilde, “Data-processing strategies for metabolomics studies,” TrAC-Trends Anal. Chem., vol. 30, no. 10, pp. 1685–1698, 2011.

[13] M. Bastian, S. Heymann, and M. Jacomy, “Gephi: An Open Source Software for Exploring and Manipulating Networks Visualization and Exploration of Large Graphs.”

[14] Christina Steppeler, “Effects of meat and meat components on intestinal carcinogenesis in the A/J Min/+ mouse model,” NMBU - Norwegian University of Life Sciences, 2017.

[15] J. G. Caporaso, J. Kuczynski, J. Stombaugh, K. Bittinger, F. D. Bushman, E. K. Costello, N. Fierer, A. G. Peña, J. K. Goodrich, J. I. Gordon, G. A. Huttley, S. T. Kelley, D. Knights, J. E. Koenig, R. E. Ley, C. A. Lozupone, D. McDonald, B. D. Muegge, M. Pirrung, J. Reeder, J. R. Sevinsky, P. J. Turnbaugh, W. A. Walters, J. Widmann, T. Yatsunenko, J. Zaneveld, and R. Knight, “QIIME allows analysis of high-throughput community sequencing data,” Nat. Methods, vol. 7, no. 5, pp. 335–336, May 2010.

[16] R. Bro and A. K. Smilde, “Centering and scaling in component analysis,” J. Chemom., vol. 17, no. 1, pp. 16–33, Jan. 2003.

[17] R. Bro, “PARAFAC. Tutorial and applications,” Chemom. Intell. Lab. Syst., vol. 38, no. 2, pp. 149–171, 1997.

[18] G. Tomasi and R. Bro, “PARAFAC and missing values,” Chemom. Intell. Lab. Syst., vol. 75, no. 2, pp. 163–180, 2005.

[19] C. A. Andersson and R. Bro, “The N-way Toolbox for MATLAB,” Chemom. Intell. Lab. Syst., vol. 52, pp. 1–4, 2000.

[20] A. K. Smilde, J. J. Jansen, H. C. J. Hoefsloot, R. J. a N. Lamers, J. van der Greef, and M. E. Timmerman, “ANOVA-simultaneous component analysis (ASCA): A new tool for analyzing designed metabolomics data,” Bioinformatics, vol. 21, no. 13, pp. 3043–3048, Jul. 2005.

[21] A. El Ghaziri, E. M. Qannari, T. Moyon, and M.-C. Alexandre-Gouabau, “AoV-PLS: a new method for the analysis of multivariate data depending on several factors,” Electronic Journal of Applied Statistical Analysis, vol. 8, no. 2. pp. 214–235, 2015.

[22] Ø. Langsrud, “50–50 multivariate analysis of variance for collinear responses,” J. R. Stat. Soc. Ser. D, vol. 51, no. 3, pp. 305–317, 2002.

[23] “Nofima modelling downloads.” [Online]. Available: http://nofimamodeling.org/software-downloads-list/.

[24] R. Bro, “Multiway calibration. Multilinear PLS,” J. Chemom., vol. 10, no. 1, pp. 47–61, Jan. 1996.

[25] J. A. Westerhuis, T. Kourti, and J. F. MacGregor, “Analysis of multiblock and hierarchical PCA and PLS models,” J. Chemom., vol. 12, no. 5, pp. 301–321, 1998.

[26] T. Næs, O. Tomic, N. K. Afseth, V. Segtnan, and I. Måge, “Multi-block regression based on combinations of orthogonalisation, PLS-regression and canonical correlation analysis,” Chemom. Intell. Lab. Syst., vol. 124, pp. 32–42, 2013.

[27] B. Moen, A. Oust, Ø. Langsrud, N. Dorrell, G. L. Marsden, J. Hinds, A. Kohler, B. W. Wren, and K. Rudi, “Explorative multifactor approach for investigating global survival mechanisms of Campylobacter jejuni under environmental conditions.,” Appl. Environ. Microbiol., vol. 71, no. 4, pp. 2086–94, Apr. 2005.

[28] Ø. Langsrud, “Rotation tests,” Stat. Comput., vol. 15, no. 1, pp. 53–60, Jan. 2005.

[29] T. Mehmood, K. H. Liland, L. Snipen, and S. Sæbø, “A review of variable selection methods in Partial Least Squares Regression,” Chemom. Intell. Lab. Syst., vol. 118, pp. 62–69, Aug. 2012.

[30] S. Wold, E. Johansson, and M. Cocci, “PLS: Partial Least Squares Projections to Latent Structures,” in 3D QSAR in Drug Design: Volume 1, 1993, pp. 523–550.

[31] L. Eriksson, T. Byrne, E. Johansson, J. Trygg, and C. Wikström, Multi-and Megavariate Data Analysis, 3rd ed. Umetrics Academy, 2013.

[32] S. Favilla, C. Durante, M. L. Vigni, and M. Cocchi, “Assessing feature relevance in NPLS models by VIP,” Chemom. Intell. Lab. Syst., vol. 129, pp. 76–86, 2013.

[33] P. J. Rousseeuw, “Silhouettes: A graphical aid to the interpretation and validation of cluster analysis,” J. Comput.Appl. Math., vol. 20, pp. 53–65, Nov. 1987.

[34] M. Halkidi, Y. Batistakis, and M. Vazirgiannis, “On Clustering Validation Techniques,” J. Intell.Inf. Syst., vol. 17, no. 2/3, pp. 107–145, 2001.

[35] K. V. Mardia, J. T. Kent, and J. M. Bibby, Multivariate analysis. Academic Press, 1979.

[36] M. Jacomy, T. Venturini, S. Heymann, and M. Bastian, “ForceAtlas2, a Continuous Graph Layout Algorithm for Handy Network Visualization Designed for the Gephi Software,” PLoS One, vol. 9, no. 6, p. e98679, Jun. 2014.

[37] M. Nei and S. Kumar, “Phylogenetic inference: Distance methods,” in Molecular evolution and phylogenetics, Oxford University Press, 2000, p. 333.

[38] K.-P. Ossenkopp, L. Sorenson, and D. S. Mazmanian, “Factor analysis of open-field behavior in the rat (Rattus norvegicus): application of the three-way PARAFAC model to a longitudinal data set,” Behav. Processes, vol. 31, no. 2–3, pp. 129–144, 1994.

[39] T. Miettinen, T. J. Hurse, M. A. Connor, S.-P. Reinikainen, and P. Minkkinen, “Multivariate monitoring of a biological wastewater treatment process: a case study at Melbourne Water’s Western Treatment Plant,” Chemom. Intell.Lab. Syst., vol. 73, no. 1, pp. 131–138, 2004.

[40] A. Niazi, J. Ghasemi, and A. Yazdanipour, “PARAFAC Decomposition of Three-Way Kinetic-Spectrophotometric Spectral Matrices Based on Phosphomolymbdenum Blue Complex Chemistry for Nitrite Determination in Water and Meat Samples,” Anal. Lett., vol. 38, no. 14, pp. 2377–2392, Nov. 2005.

[41] P. M. Kroonenberg and R. A. Harshman, “ANALYSING THREE-WAY PROFILE DATA USING THE PARAFAC AND TUCKER3 MODELS ILLUSTRATED WITH VIEWS ON PARENTING,” Appl. Multivar.Res., vol. 13, no. 1, pp. 5–41, 2009.

[42] M. Thiel, B. Féraud, and B. Govaerts, “ASCA+ and APCA+: Extensions of ASCA and APCA in the analysis of unbalanced multifactorial designs,” J. Chemom., vol. 31, no. 6, p. e2895, Jun. 2017.

